# Transcriptomic response of *Cinachyrella* cf. *cavernosa* sponges to spatial competition

**DOI:** 10.1101/2021.07.05.451097

**Authors:** Aabha Deshpande, Ramón E. Rivera-Vicéns, Narsinh L. Thakur, Gert Wörheide

## Abstract

Spatial competition in the intertidal zones drives the community structure in marine benthic habitats. Organisms inhabiting these areas not only need to withstand fluctuations of temperature, water level, pH, and salinity, but also need to compete for the best available space. Sponges are key members of the intertidal zones, and their life history processes (e.g. growth, reproduction, and regeneration) are affected by competition. Here we used transcriptomics to investigate the effects of interspecific competition between the tetillid sponge *Cinachyrella* cf. *cavernosa*, the zoantharid *Zoanthus sansibaricus*, and the macroalgae *Dictyota ciliolata*. The analysis of differentially expressed genes showed that *Z. sansibaricus* was the most stressful competitor to *C*. cf. *cavernosa*, which showed an increased rate of cellular respiration under stress of competition. Similarly, an up-regulation of energy metabolism, lipid metabolism, and the heat-shock protein (HSP) 70 was also observed along with an indication of a viral infection and decreased ability to synthesise protein. A down-regulation of purine and pyrimidine metabolism indicated reduction in physiological activities of the competing sponges. Moreover, a putative case of possible kleptocnidism, not previously reported in *Cinachyrella* cf. *cavernosa* was also observed. This study opens the door for more detailed investigations of marine organisms competing for spatial resources using transcriptome data.

## 1. Introduction

Intertidal zones, or the area of the coastline between low and high tides, are an ecosystem full of biodiversity and of great interest for many ecological studies (Lubchenco & Menge, 1978; Song et al., 2017; Tomanek & Helmuth, 2002). These zones are characterised by harsh conditions related to dramatic changes in water-level, temperature, and salinity (Satyam & Thiruchitrambalam, 2018). One of the main limiting factors shared by all benthic organisms in the intertidal zones is space. Thus, competition among organisms in the intertidal area shapes community structure and controls biodiversity (Paine, 1984; Worm & Karez, 2002).

Some of the mechanisms that organisms use to mitigate competition for space include crowding, whiplashing, overgrowing, surface-sloughing, undercutting, crushing, overshadowing, digesting, and poisoning (Bakus, Targett, & Schulte, 1986). Most previous studies mainly focused on algae and sessile invertebrates, such as mussels and barnacles (e.g., Quinn, 1982; Worm & Karez, 2002). However, other organisms, such as corals and sponges, have not been considered in detail so far, although sponges, for example, are key members of the intertidal zones that compete for space with other inhabitants. In the case of sponges of coral reefs, chemically mediated interactions among sponges and other organisms have been documented (e.g., Wulff, 2006) but no such reports have come out of intertidal zones.

Opportunistic competitors like macroalgae may force sponges to adapt. Palumbi (1985) showed that macroalgae *Corallina vancouveriensis* out competed sponge *Halichondria panacea* in the rocky intertidal area of Alaska. Singh and Thakur (2016, 2018) reported that the sponge *Cinachyrella* cf. *cavernosa* (Lamarck, 1815) competes for space with the zoantharid anthozoan *Zoanthus sansibaricus* in the intertidal zones. Furthermore, recent field observations have identified the macroalgae *Dictyota ciliolata* to be abundant in these areas (Pereira & Almeida, 2014), likely becoming one of the main competitors for space. Hence, we decided to investigate the effect of these two space competitors on the sponge *Cinachyrella* cf. *cavernosa* at transcriptomic level. We did so, because differential gene expression analysis allowed us to obtain more comprehensive insights into the sponge’s reaction to competition, including its metabolism, cellular activity, and molecular functioning. An important question was whether *C*. cf. *cavernosa* up-regulated any pathways of secondary metabolites biosynthesis to mitigate competition.

Our results suggest that *Zoanthus sansibaricus* is the most stressful competitor to *Cinachyrella* cf. *cavernosa* in the intertidal zones, as the expression of several key genes changed and a viral infection was observed. However, we did not observe any significant up-regulation in biosynthesis of secondary metabolites.

## 2. Methods

### 2.1 Study site and Sample collection

Samples were collected from the rocky beach of Anjuna, Goa, India (15°34’24”N - 73°44’25”E, subtropical zone) in March 2019. At this site, clear zonation can be seen where only *Zoanthus sansibaricus* (Phylum Cnidaria, class Anthozoa) is found at the lower intertidal area. Macroalgae are more abundant towards the upper intertidal area as compared to the lower intertidal area. However, the sponge *Cinachyrella* cf. *cavernosa* can be found along the entire stretch from lower to upper intertidal area. Samples were divided as: a) sponges without competition (Fig 1A, referred hereafter as S) from the mid intertidal area; b) sponges in competition with macroalgae *D. ciliolata* (Fig 1B, referred hereafter as SA) from the upper intertidal area; c) sponges in competition with *Z. sansibaricus* (Fig 1C, referred hereafter as SZ) from the lower intertidal area. Six replicates of each sample were collected, cut with a sterile scalpel, kept in sterile vials, and transferred on ice before flash-freezing in liquid nitrogen. Samples were stored at −80°C until RNA extraction.

**Figure 1:**
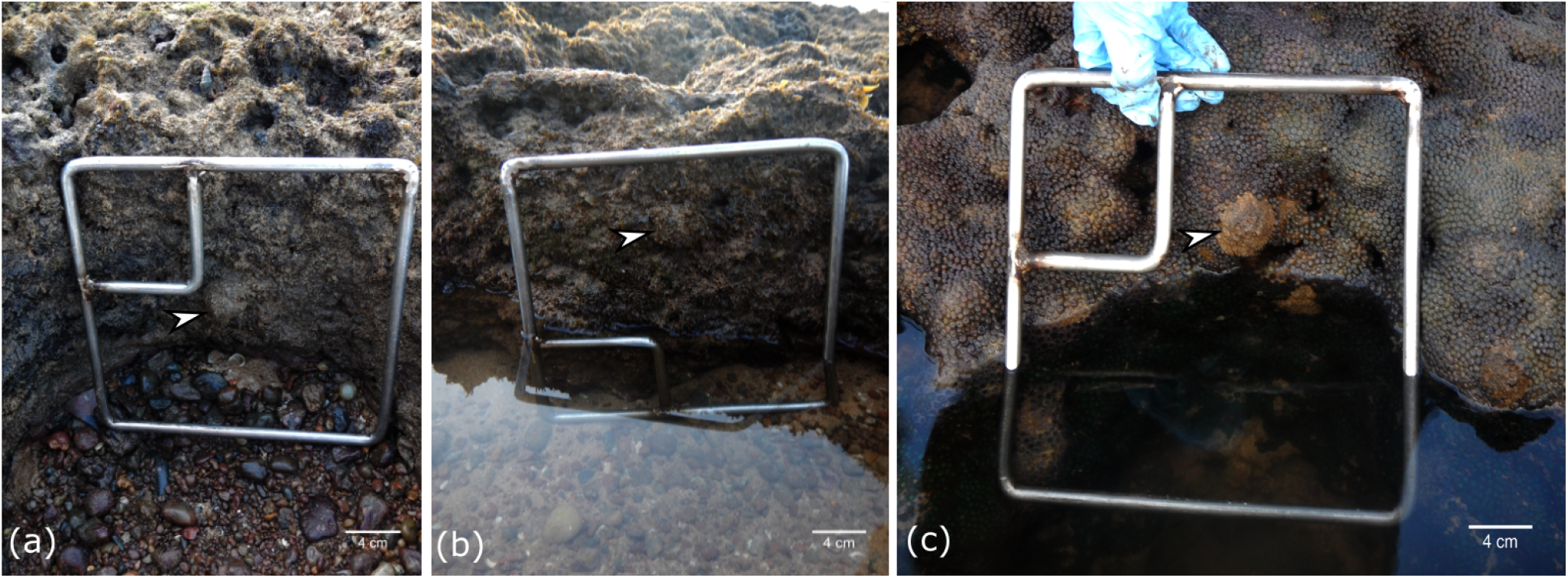
Sponge and its neighbours. Sponge *Cinachyrella* cf *cavernosa* in each photo is indicated by a white arrowhead. (a) sponge with no competitors; (b) Sponge with macroalgae *Dictyota ciliolata* as competitor; (c) Sponge with soft coral *Zoanthus sansibaricus* as competitor.

### 2.2 RNA extraction, library preparation, and sequencing

Total RNA was isolated using the conventional TRIzol method. The quality and quantity of RNA from each sample were checked on 1% denaturing RNA agarose gel and NanoDrop, respectively. Three samples from each group were used for sequencing. The RNA-Seq paired-end sequencing libraries were prepared using Illumina TruSeq stranded mRNA sample prep kit. Libraries were analysed in 4200 TapeStation systems (Agilent Biotechnologies) using high sensitivity D5000 Screen tape. The mean peak size was obtained along with Qubit concentration of libraries. Illumina paired-end libraries were sequenced using a NextSeq500 with a read length of 75bp. RNA extraction and transcriptome sequencing was carried out by Eurofins Genomics India Pvt Ltd, Bangalore, India. Sequences were deposited in the Annotare archive (https://www.ebi.ac.uk/arrayexpress/experiments/E-MTAB-9052/) of the European Bioinformatics Institute under accession number E-MTAB-9052.

### 2.3 Assembly and annotation

For the generation of the de novo transcriptome assemblies the reads were processed using TransPi (Rivera-Vicéns, García-Escudero, Conci, Eitel, & Wörheide, 2021). Briefly, reads were assessed for quality, normalized, and assembled using a combination of different assemblers and different k-mer lengths. All these assemblies were used as input for a reduction step. This step produced a non-redundant final transcriptome assembly per individual libraries (i.e. S, SA, SZ). A reference transcriptome of *Cinachyrella* cf. *cavernosa* was created by combining all assemblies obtained from the reduction steps explained above and applying TransPi with a final reduction step. The annotation was also performed using TransPi (Rivera-Vicéns et al., 2021). It employed various databases such as SwissProt, PFAM, and a custom UniProt database (i.e., all metazoan proteins) to create a final report including information on gene ontology (GO), eggNOG, KEGG, PFAM, proteins similarities to SwissProt and UniProt, among others. Annotation terms of eggNOG and KEGG were visualized with iPath3.0 (Darzi, Letunic, Bork, & Yamada, 2018). The transcriptome assembly is available in European Nucleotide Archives with Study accession number PRJEB39339.

### 2.4 Differential expression analysis

Using the reference assembly described above, transcript abundance and the genes count matrix for all individuals were created using RSEM-Bowtie2 (v0.8.0; Li & Dewey, 2011). DESeq2 (v1.24.0; Love, Huber, & Anders, 2014) was used to select the differentially expressed genes with a p-value cut-off 0.01 and minimum log2 (fold change) of two. The analyses consisted of various test scenarios: a) control vs. competitors (algae + soft coral i.e., all samples; SvAll); b) control vs. algae as competitor (SvSA); c) control vs. soft coral as competitor (SvSZ); d) algae as competitor vs. soft coral as competitor (SAvSZ). Gene ontology enrichment was performed with topGO (p <0.01) (Alexa and Rahnenfuhrer, 2016) using the differentially expressed genes and the GO terms file obtained from the annotation.

## 3 Results

### 3.1 Reference assembly

High quality reads were obtained for all nine samples (three samples per group) and did not require trimming (Table 1). No over-represented sequences or adaptors were found in the reads. The average number of base pairs for the nine samples was 31,608,536. The GC content was consistent, with an average of 49.7% across all samples. The reference assembly had a total of 117,382 transcripts, N50 of 1,076, and GC content of 48.5% (Table 2). BUSCO scores for the reference assembly and individuals are presented on Figure 2a. Reference assembly had 92.22% of the metazoan dataset of BUSCO (Figure 2a). Total BUSCO genes presence found in the reference assembly was 96.5%. Individual assemblies (i.e. each condition, viz., S, SA, SZ) had BUSCO scores between 85.89% – 91.41% where most of these genes were found to be single copy (71.81%). This difference is not much significant since we performed multiple rounds of reductions steps to get our final reference transcriptome (see Methods), and combining all individual assemblies did not have a major effect in terms of duplicated sequences. Moreover, our reference transcriptome covers most of the sequencing reads obtaining a mapping percentage of 81.76% (after reduction of duplicates) This represents an increment of around 3.95% on the mapping rate when compared to the individual assemblies.

**Figure 2:**
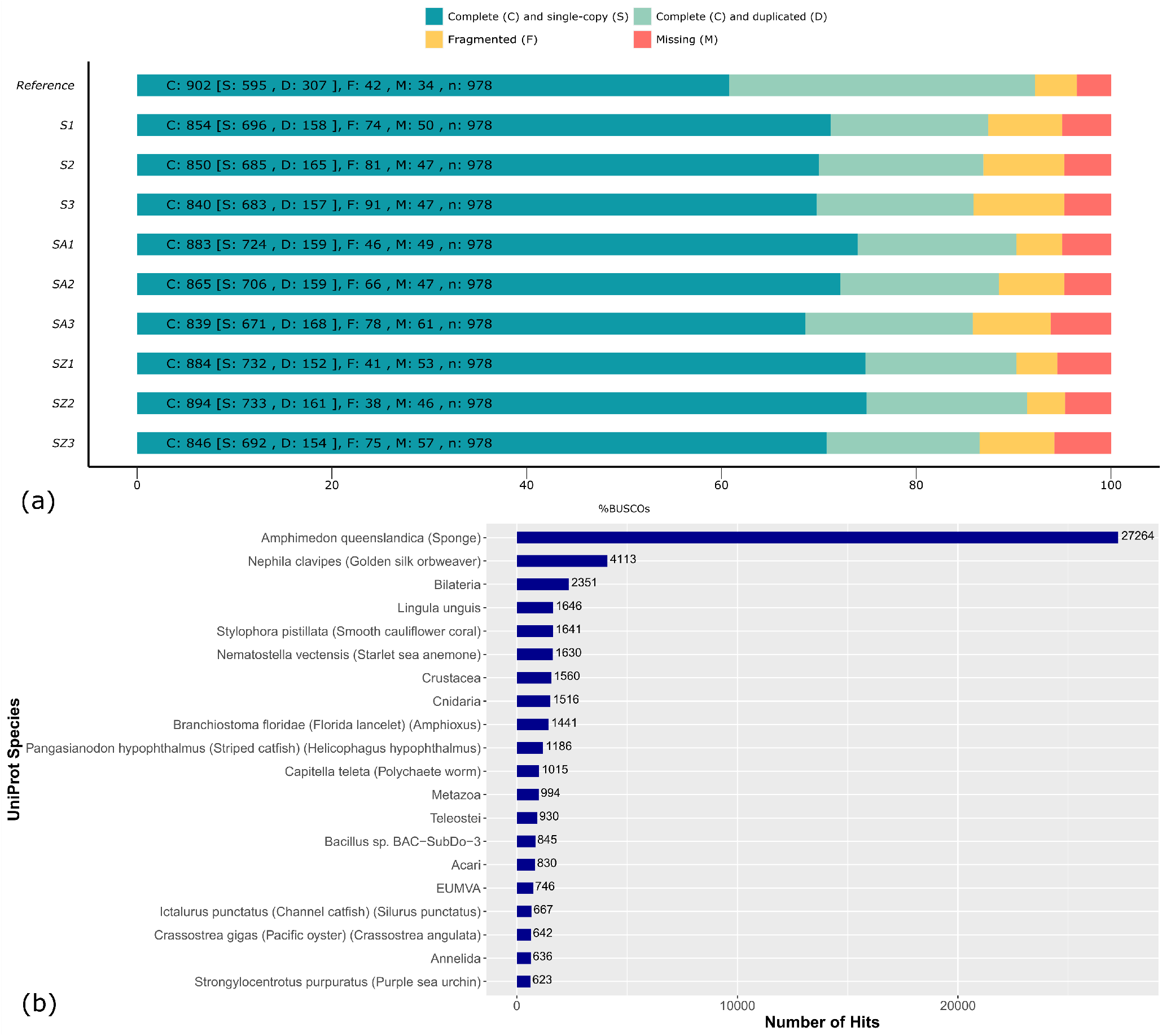
Completeness and annotation of the transcriptome assembly (a) Comparison between the completeness of the reference assembly and the individual assemblies; (b) Annotation of the reference assembly shown as the hits against a particular organism.

**Table 1:** Quality statistics of the raw reads

| Sample | No. of base pairs | GC% | No. of transcripts | Transcripts>1000 | N50 |
| --- | --- | --- | --- | --- | --- |
| S1 | 32,839,566 | 49.9 | 34,345 | 10,963 | 1,558 |
| S2 | 32,289,462 | 49.1 | 37,116 | 10,567 | 1,417 |
| S3 | 31,913,009 | 49 | 37,702 | 10,228 | 1,339 |
| SA1 | 31,126,654 | 49.3 | 34,195 | 10,791 | 1,502 |
| SA2 | 29,674,960 | 49.6 | 31,985 | 10,237 | 1,443 |
| SA3 | 31,640,674 | 51.8 | 31,601 | 10,610 | 1,552 |
| SZ1 | 32,449,014 | 48.6 | 34,423 | 11,175 | 1,570 |
| SZ2 | 31,421,326 | 49.1 | 32,738 | 10,785 | 1,597 |
| SZ3 | 31,122,159 | 50.9 | 34,788 | 10,151 | 1,379 |

**Table 2:**
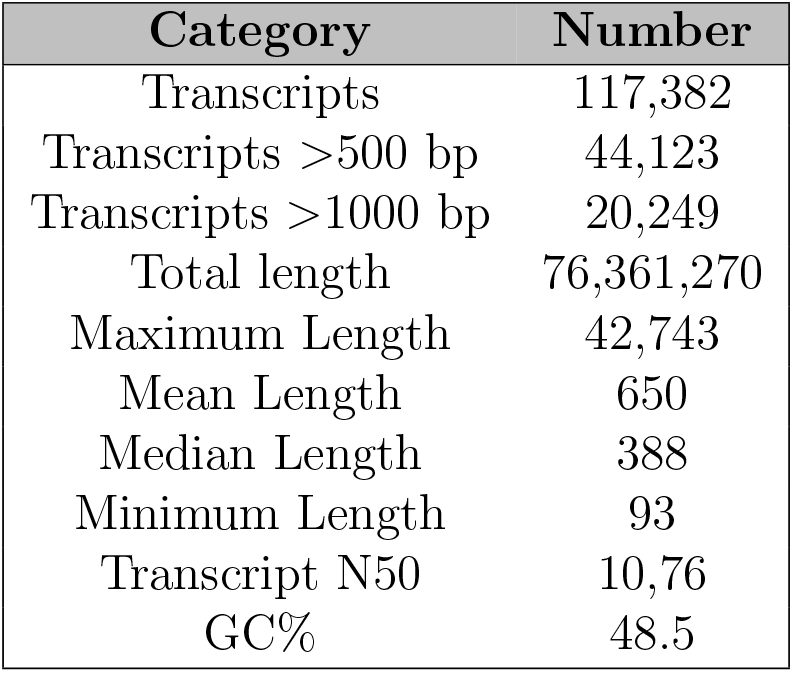
Quality statistics of the reference assembly

### 3.2 Annotation

In the automated annotation executed by TransPi, a total of 65,089 ORFs (using homology retention criteria) (Table 3) for the reference transcriptome were obtained. The majority of these ORFs (56,719) had hits to either PFAM and/or UniProt (retention criteria). SwissProt database search had 35,689 hits with BLASTX and 29,246 with BLASTP. From these hits, information on Gene Ontology, KEGG and eggNOG was obtained. Protein from the SwissProt database with most hits (n=119) was for the Major basic nuclear protein 1 (HCC2_CRYCO) from *Crypthecodinium cohnii*. In the second BLAST search against all metazoan UniProt proteins a total of 45,363 and 37,872 hits were obtained for BLASTX and BLASTP, respectively. Protein with most hits (n=102) was to the Eukaryotic elongation factor 1 alpha (A1L2K7_XENLA) from *Xenopus laevis*. The annotation analysis also revealed that the majority of our sequences had similarities with sequences from *Amphimedon queenslandica*(n=27,264). Other hits were to cnidarians (*Stylophora pistillata, Nematostella vectensis*), crustaceans, bilateria, brachiopod, and metazoans proteins (Figure 2b). However, since the most of our proteins were matched to the sponge *A. queenslandica* we did not perform extra steps to get rid of these sequences.

**Table 3:** Summary of gene ontology of the reference transcriptome

| Sample | eggNOG and COG | GO | KEGG | PFAM | BLAST | ORFs | %GC |
| --- | --- | --- | --- | --- | --- | --- | --- |
| S1 | 10,147 | 12,322 | 8,871 | 63 | 16,778 | 25,807 | 49.92 |
| S2 | 10,474 | 13,063 | 9,332 | 49 | 17,285 | 26,084 | 49.14 |
| S3 | 11,544 | 14,403 | 10,301 | 56 | 18,997 | 28,292 | 48.95 |
| SA1 | 10,307 | 13,851 | 8,972 | 15,085 | 18,580 | 24,468 | 48.63 |
| SA2 | 9,707 | 11,725 | 8,531 | 68 | 16,127 | 24,643 | 49.15 |
| SA3 | 11,548 | 14,860 | 10,223 | 16,705 | 19,894 | 26,268 | 50.9 |
| SZ1 | 10,286 | 13,849 | 8,925 | 15,134 | 18,335 | 24,587 | 49.28 |
| SZ2 | 9,411 | 11,591 | 8,165 | 55 | 15,984 | 23,451 | 49.56 |
| SZ3 | 10,051 | 12,480 | 8,839 | 75 | 17,166 | 26,946 | 51.79 |
| all | 26,277 | 37,910 | 24,135 | 38,178 | 46,663 | 65,089 | 48.5 |

Gene ontology of the *Cinachyrella* cf. *cavernosa* reference transcriptome is presented on Figure 2b. Overall 91,202, 143,979 and 105,008 hits were obtained for Molecular Functions (MF), Biological Processes (BP) and Cellular Components (CC), respectively. Ontology classification with most hits were for ATP binding (n=6,186, MF), Translation (n=2,895, BP), and Cytoplasm (n=9,996, CC). Annotated transcripts were visualized using eggNOG and KEGG terms. Summary of the complete annotation results is presented in Table 3.

### 3.3 Differential Expression

Differential expression analysis of control vs sponges with all competitors (algae + soft coral) using DESeq2 showed 614 up-regulated genes and 114 down-regulated genes. This difference was the largest among all the DEG analyses we performed (Table 4). From the DEGs, 63 up-regulated genes and eight down-regulated genes had annotations to SwissProt. A total of 71 DEGs out of 737 were annotated (9.6%). Heatmaps summarizing all these analyses are presented in Figure 3 (A=up, B=down).

**Figure 3:**
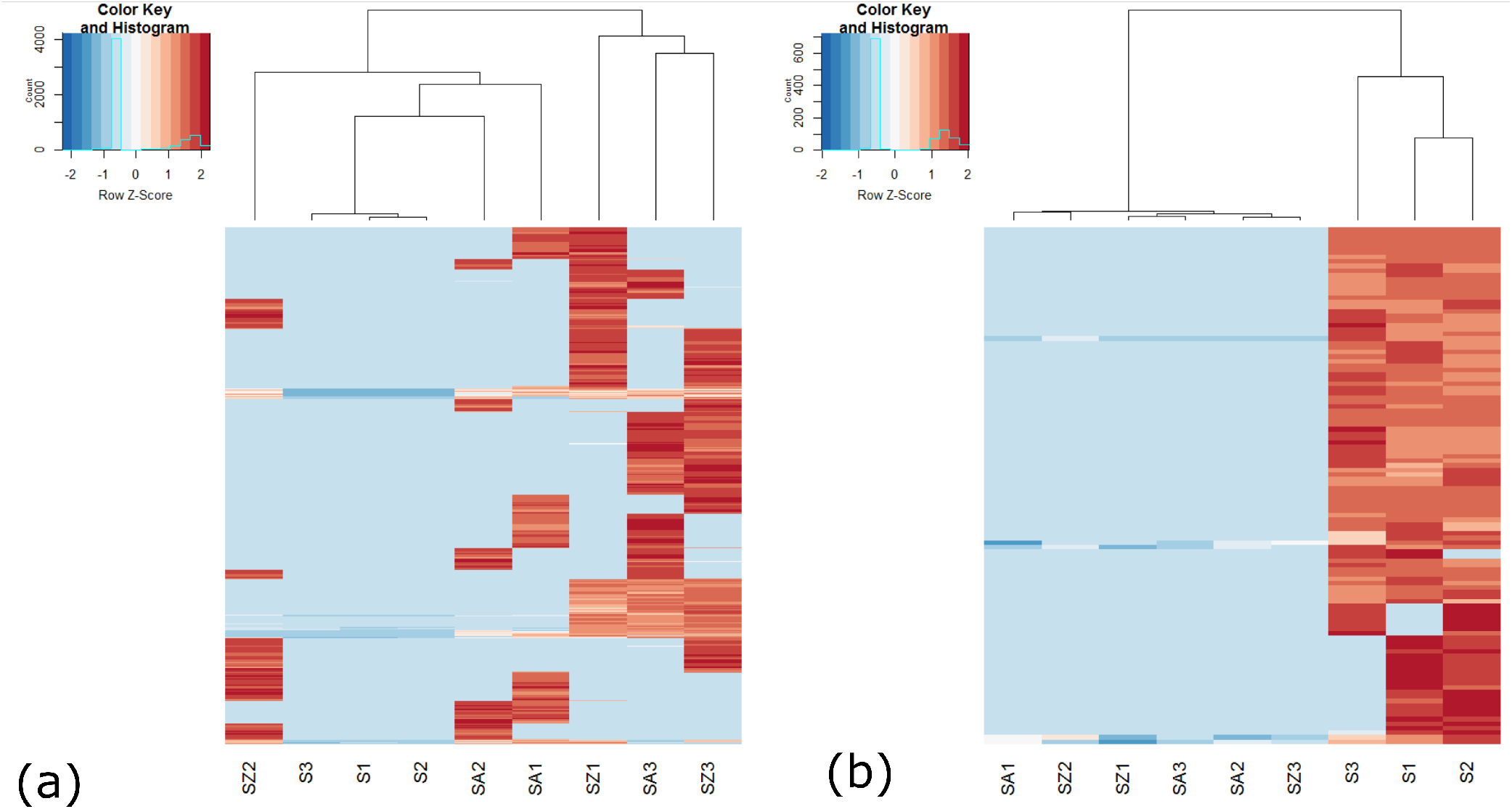
Heatmaps showing difference in DEGs among all samples (A) Up-regulated DEGs for SvAll; (B) Down-regulated DEGs for SvAll

**Table 4:** Summary of results of differential expression analysis

| Condition | Up-regulated genes | Down-regulated genes |
| --- | --- | --- |
| SvAll | 614 | 114 |
| SvSA | 16 | 133 |
| SvSZ | 32 | 302 |
| SAvSZ | 58 | 145 |

To further analyze expression patterns we also performed DEGs analyses for the SvSA, SvSZ, SAvSZ samples (Table 4; Supplementary information Table S1). More number of down-regulated genes were obtained as compared to up-regulated genes. This pattern is exactly opposite to those observed in control vs all competitors samples. Differential expression analysis for the sponge versus the soft coral (SvSZ) resulted in 32 and 302 up-regulated and down-regulated genes, respectively. The analysis of sponge vs algae (SvSA) showed a total of 16 up-regulated genes and 133 down-regulated genes. The results are much lower than the DEGs obtained for sponge vs soft coral (SvSZ). For the samples consisting of sponges with algae and sponge with soft coral (SAvSZ), 58 DEGs were up-regulated and 145 DEGs were down-regulated (Table 4).

### 3.4 Enrichment analysis

From the up-regulated genes of sponges without competitors i.e. control vs sponges with all competitors (algae + soft coral), a total of 25 terms were enriched in the CC (Cellular Component) classification, 21 in MF (Molecular Function) and 76 in the BP (Biological Process) (Figure 4a, b, c). Enrichment of BP terms were associated with cellular respiration and oxidative phosphorylation (Figure 4a). MF terms were associated with electron transfer activity and NADH dehydrogenase activity (Figure 4c). Among the enriched terms for CC, we found terms for respiratory chain and various mitochondrial related components (Figure 4b). These enriched terms can also be visualized by mapping the DEGs to the KEGG pathways where up-regulation occurs for the oxidative phosphorylation paths (Figure 5, blue color). Among the up-regulated DEGs, many collagen, actin, and tubulin related genes which are involved in cytoskeleton stabilization and intra-cellular transport were observed. Genes related to myogenesis and muscle contraction in other animals were also found. Up-regulation of fatty acid metabolism and expression of HSP70, which are known responses for stress (Benette, Bell, Davy, Webster, & Francis, 2017; Lopez-Legentil, Song, Macmurray, & Pawlik, 2008) were observed. Sponge immune response related genes (e.g., TRAF3, ILF2) were seen to be up-regulated. Lastly, an up-regulation and down-regulation of different components related to cellular respiration was observed. For the down-regulated genes, 22, 26, 34 terms were enriched for CC, MF, and BP classifications, respectively (Figure 4d, e, f). Enrichment for BP terms are associated with oxidative phosphorylation (Figure 4d). For MF terms, enrichment was associated with cytochrome c oxidase activity (Figure 4f), whereas electron transfer activity and NADH dehydrogenase activity were the CC terms (Figure 4e).

**Figure 4:**
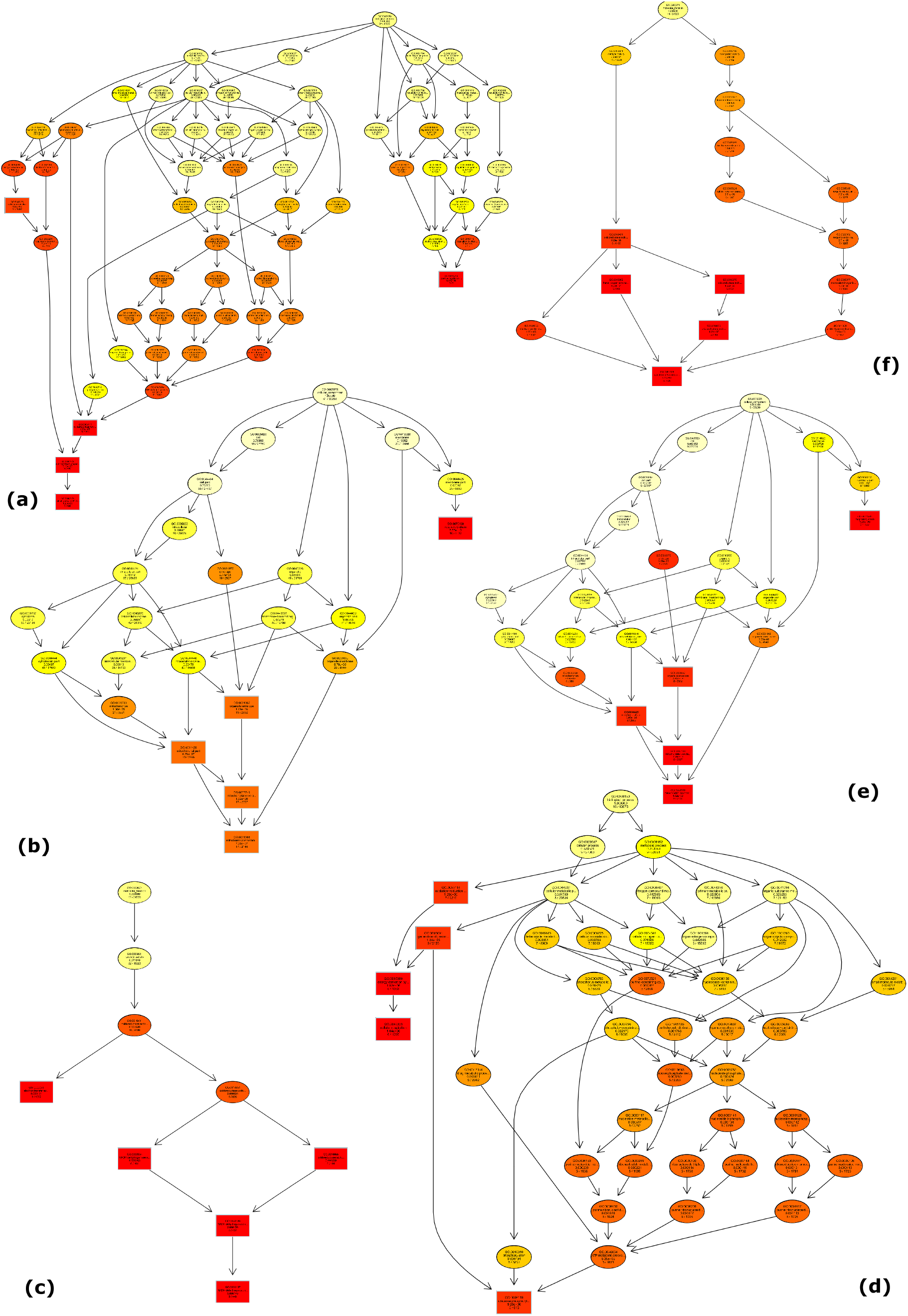
Enrichment of gene ontology terms of DEGs from SvAll (a) Up-regulated biological processes (BP) terms; (b) Up-regulated cellular components (CC) terms; (c)Up-regulated molecular functions (MF) terms; (d) Down-regulated BP terms; (e) Down-regulated CC terms; (f) Down-regulated MF terms

**Figure 5:**
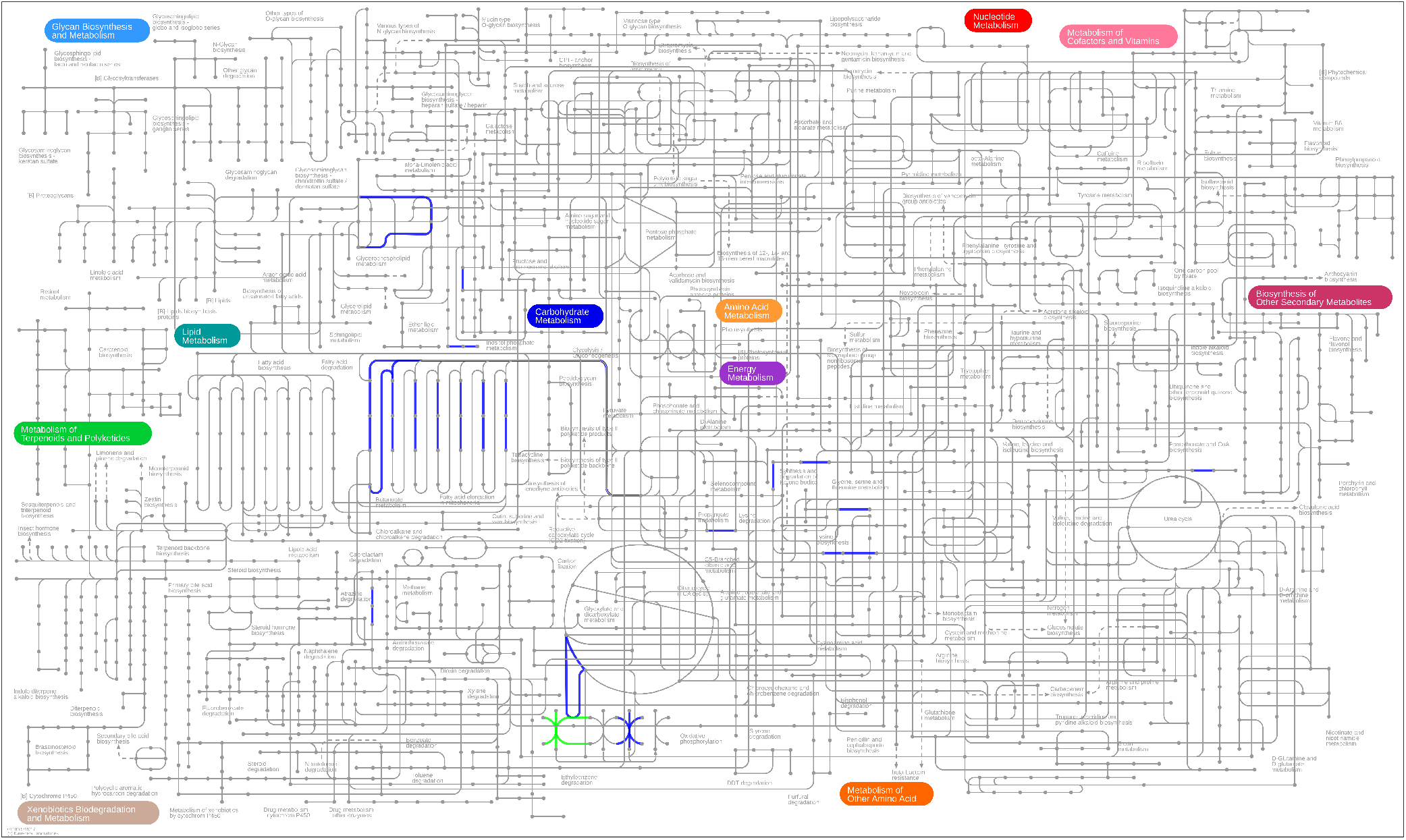
Map showing up-regulated (blue) and down-regulated (green) KEGG pathways. The interactive version can be accessed at [here](https://pathways.embl.de/ipath3.cgi?s=ncur0N9NIgemk9LhUb7&map=metabolic)

#### Sponges without competitors vs sponges with soft coral *Zoanthus sansibaricus* as the competitor

Enrichment of up-regulated genes showed a significant enrichment of Rab GTPase binding activity and increased GTPase activity for the MF terms. In the BP terms, an enrichment for positive regulation of viral proteins by virus was identified. An enrichment of CC terms related to positive regulation of endoplasmic reticulum to Golgi vesicle-mediated transport was also observed. SvSZ samples also showed CC terms enrichment in the endoplasmic reticulum and Golgi intermediate compartment and the component of Golgi membrane consisting of gene products and protein complexes. For the down-regulated genes, oxidative phosphorylation and purine and pyrimidine metabolism were enriched for the BP terms. For the MF terms, enrichment was observed for the electron transfer activity and cytochrome c oxidase activity. CC terms enriched were for mitochondrial inner membrane and organelle inner membrane (Table S1 a and b).

#### Sponges without competitors vs sponges with macroalgae *Dictyota ciliolata* as the competitor

Enrichment of up-regulated genes for the MF category was observed for copper ion binding and cytochrome c oxidase activity. An enrichment of the electron transport chain and the generation of the precursor of metabolites and energy was observed for the BP terms. In the case of CC terms an enrichment was observed for terms associated with nematocysts and cell cortex. For the down-regulated genes, the enrichment of BP terms was observed for aerobic respiration and oxidative phosphorylation. For the MF terms, the enrichment analysis showed that cytochrome c oxidase activity and heme copper oxidase activity were enriched. Last, the enriched CC terms were respiratory chain complex IV and cytochrome complex (Table S1 c and d).

#### Sponges with macroalgae *Dictyota ciliolata* as the competitor vs sponges with soft coral *Zoanthus sansibaricus* as the competitor

Enriched up-regulated BP terms were for aerobic respiration and oxidative phosphorylation. For the CC terms, enriched terms were cytochrome c oxidase activity and heme copper oxidase activity. CC terms for respiratory chain complex IV and cytochrome complex were also enriched. MF terms were enriched for cytochrome-c oxidase activity and heme binding activity. For the down-regulated genes, translation and peptide biosynthetic processes were enriched for the BP terms. In the case of MF terms, enrichment was found for structural constituents of ribosome and structural molecule activity. CC terms enriched were for ribosome and ribonucleoprotein complex (Table S1 e and f).

## 4 Discussion

Sponges are key members of the intertidal zones. They compete for space with organisms like corals and algae as this controls the community composition and biodiversity of the area. Most studies have focused on sessile invertebrates (e.g., Quinn, 1982; Worm & Karez, 2002). Few have described the chemical interaction of sponges while competing (Wulff, 2006). However, none of the reports included the intertidal zones. Thus, an investigation was conducted by employing transcriptomic techniques to understand the space competition in the intertidal zone between the sponge *Cinachyrella* cf. *cavernosa* and the zoantharid *Zoanthus sansibaricus* and macroalgae *Dictyota ciliolata*.

The assembled *Cinachyrella* cf. *cavernosa* reference transcriptome presents a significant improvement over previous *C*. cf. *cavernosa* transcriptome (Deshpande & Thakur, 2020). Furthermore, our BUSCO results (i.e., 96.55%) surpass all published sponge transcriptomes at the moment, which are found in the range of 69-94% (González-Aravena et al., 2019; Kenny et al., 2018; Kenny, Plese, Riesgo, & Itskovic, 2019; Pita, Hoeppner, Ribes, & Hentschel, 2018). The majority of the complete BUSCO genes from our reference assembly were single copy, indicating a non-redundant consensus assembly (Simão, Waterhouse, Ioannidis, Kriventseva, & Zdobnov, 2015; Waterhouse, Zdobnov, & Kriventseva, 2011; Table2, Figure 2a). Annotation showed high similarity to sponge *A. queenslandica*. Protein hits to other species could indicate a possible source of contamination. However, it also presents the problem of under annotation that non-model species, like porifera (Renard, Leys, Wörheide, & Borichiellini, 2018), currently have, where a protein from a distant, unrelated organism is a match in the absence of a better candidate. For example, various hits from the custom metazoan uniprot dataset were assigned to the spider *Nephila clavipes*, but at the same time this sequences were assigned to different classes of bacteria (e.g., Alphaproteobacteria, Betaproteobacteria) in the SwissProt database. These could then represent sequences from the bacterial symbionts and not from contamination sources.

### 4.1 Differential expression and enrichment

#### Sponge without competitors vs Sponge with all the competitors

Comparing multiple competitors at the same time detected more changes from all of the DEG analyses that were conducted. Up-regulation of genes was approximately five times higher than the down-regulated genes. Up-regulation of genes related to myogenesis and muscle contraction was observed in the enrichment analysis. Although sponges lack muscles they are known to exhibit coordinated behaviour such as contraction (Nickel, 2010). These sponges under competition could be contracting their oscules to avoid uptake of harmful particles from the water canals (Leys, 2015).

In a previous study the expression of HSP 70 in *C*. cf. *cavernosa* was shown to increase when competing with *Z. sansibaricus* (Singh & Thakur, 2018). Our analysis also showed an up-regulation of HSP70 along with fatty acid metabolism. Moreover, there is an increased energy production i.e., up-regulation of ATP synthesis and hydrolysis, and reduction in metabolic activity of the sponge in terms of down-regulation cytochrome c oxidase and NADH oxidoreductase chains (Figure 2b).

Enrichment of the up-regulated genes also indicated increased energy demand due to the stress of competition. In the KEGG pathway, paths of the lipid metabolism were up-regulated, specifically pathways of fatty acid elongation. This is in agreement with previous reports that addressed the role of lipids in stress management in sponges (Benette et al., 2017; Deshpande & Thakur, 2020). It should be noted that some of the enriched down-regulated genes were found to be enriched in the up-regulated genes as well. These could be a result of using both competitors together (algae + soft coral) in the analysis. However, the pattern is resolved when analyzing individual interactions.

#### Sponges without competitors vs sponges with soft coral *Zoanthus sansibaricus* as the competitor

Our analysis indicates that the competition with zoanthid *Z. sansibaricus* causes more changes in the sponge as compared to competition with *D. ciliolata*. Since they belong to the same trophic level, both organisms (i.e., sponge and zoanthid) are not only competing for space, but for food sources as well. Enrichment of up-regulated genes suggests opportunistic viral infections in already stressed and vulnerable sponges. Even though there is an acute lack of studies on viruses associated with marine sponges (Webster & Taylor, 2011), viral infection of sponges has been previously reported (Claverie et al., 2009). Enrichment of CC terms suggest transport and localization of molecules within or between cells, possibly as a response to the viral infection or the stress of competition. Enriched down-regulated genes suggest the reduction in metabolic activity of the sponges. It is possible that the stress of competition is responsible for this decrease. In a previous study, the adverse effect of *Z. sansibaricus* was seen on the physiological activity of the same sponge (Singh & Thakur, 2016).

#### Sponges without competitors vs sponges with macroalgae *Dictyota ciliolata* as the competitor

Results suggest that the sponge is less stressed when in presence of the algae *D. ciliolata* as compared to *Z. sansibaricus*. Enrichment of up-regulated genes are involved in the liberation of energy. All these factors indicate increased energy production to mitigate stress exerted by competitors. In the case of the enrichment of CC terms, results suggest a new case of kleptocnidism in sponges. Kleptocnidism is observed when an organism (i.e., the sponge) has acquired nematocysts to use as a defense mechanism for competitors. This phenomenon has been reported in other sponges like *Haliclona* spp. (Schellenberg et al., 2019; Russell, Degnan, Garson, & Skilleter, 2003). However, there is no other evidence of presence of nematocysts in the sponge *Cinachyrella* cf. *cavernosa*. Nevertheless, this putative case of kleptocnidism warrants further thorough investigation. For the down-regulated genes, enrichment results are in contradiction with the up-regulated genes. However, since enrichment analysis deals with pathways rather than single genes, it may not be sensitive enough to differentiate between expression levels of single genes (Mootha et al., 2003; Subramanian et al., 2005). It also could be possible that different components of the same pathway are up-regulated and down-regulated.

#### Sponges with macroalgae *Dictyota ciliolata* as the competitor vs sponges with soft coral *Zoanthus sansibaricus* as the competitor

Enriched up-regulated genes suggest more energy production in SZ samples as compared to the SA samples. The up-regulation of aerobic respiration indicates more stress and increased energy production in samples in competition with the soft coral. Hence, the soft coral is considered a more aggressive competitor than macroalgae. Results from down-regulated genes are in agreement with a previous study which reports decrease in protein synthesis ability of the sponge *C*. cf. *cavernosa* when faced with the aggressive competitor *Z. sansibaricus* (Singh & Thakur, 2016). Our results suggest that *D. ciliolata* does not harm the sponge’s protein synthesis.

To conclude, *Z. sansibaricus* exerts more pressure on *C*. cf. *cavernosa* as compared to *D. ciliolata*, possibly due to overlap of spatial and trophic competition between the sponge and *Z. sansibaricus*. Sponges under stress show a significantly enriched up-regulation of oxidative phosphorylation. An up-regulation of lipid metabolism related pathways in sponges dealing with *Z. sansibaricus* as neighbours was also observed. Indications of opportunistic viral infection in these sponges and decreased protein synthesis ability was also seen. It should be noted that a putative case of kleptocnidism was observed when analyzing the enrichment data of sponges competing with *D. ciliolata*, but more in-depth studies are needed to prove the existence of this phenomenon not previously described for the sponge *C*. cf. *cavernosa*. No significant up-regulation in biosynthetic pathways for secondary metabolites was observed in sponges under stress of competition.

## Acknowledgements

We thank the Director, CSIR-NIO for providing laboratory facilities. We are grateful to Dr Temjensangba Imchen for his help in identification of the macroalgae and to Dr Nicola Conci for his help with differential expression analysis. AD thanks Council of Scientific and Industrial Research for research fellowship and European Molecular Biology Organisation (EMBO) for a short-term fellowship to visit Ludwig-Maximilians-Universität München. AD and NLT acknowledge the Council of Scientific Research funded project ‘Ocean Finder’ (PSC0105). RERV and GW acknowledge funding from the European Union’s Horizon 2020 research and innovation programme under the Marie Skłodowska-Curie grant agreement No 764840 (ITN IGNITE).

## Data Accessibility Statement

All transcriptome sequences are deposited in the Annotare archive of the European Bioinformatics Institute under accession number E-MTAB-9052. Assemblies were submitted to the European Nucleotide Archive (ENA) under accession number PRJEB39339.

## Author Contribution

AD designed the study, procured sequences, conducted differential expression analysis and wrote the manuscript. RERV carried out assembly preparation, annotation, enrichment analysis and wrote the manuscript. GW contributed to the manuscript editing, provided computational infrastructure and funding. NLT supervised study design, sequencing and provided funding.

