## Supplementary Table 1 for "Transcriptomic response of *Cinachyrella* cf. *cavernosa* sponges to spatial competition"

### Supplementary Tables

1- Sponges without competitors vs sponges with soft coral *Zoanthus sansibaricus* as the competitor – up-regulated genes

2- Sponges without competitors vs sponges with soft coral *Zoanthus sansibaricus* as the competitor – down-regulated genes

3- Sponges without competitors vs sponges with macroalgae *Dictyota ciliolata* as the competitor – up-regulated genes

4- Sponges without competitors vs sponges with macroalgae *Dictyota ciliolata* as the competitor – down-regulated genes

5- Sponges with macroalgae *Dictyota ciliolata* as the competitor vs sponges with soft coral *Zoanthus sansibaricus* as the competitor – up-regulated genes

6- Sponges with macroalgae *Dictyota ciliolata* as the competitor vs sponges with soft coral *Zoanthus sansibaricus* as the competitor – down-regulated genes

**Table 1:** Sponges without competitors vs sponges with soft coral *Zoanthus sansibaricus* as the competitor – up-regulated genes

| GO.ID | Term | Annotated | Significant | Expected | classicFisher |
|---|---|---|---|---|---|
| **Biological Processes** | | | | | |
| GO:0046726 | positive regulation by virus of viral protein levels in host cells | 3 | 2 | 0 | 6.40E-08 |
| GO:1902953 | positive regulation of ER to Golgi vesicle-mediated transport | 4 | 2 | 0 | 1.30E-07 |
| GO:0046719 | regulation by virus of viral protein levels in host cell | 5 | 2 | 0 | 2.10E-07 |
| GO:0001675 | acrosome assembly | 9 | 2 | 0 | 7.70E-07 |
| GO:0007030 | Golgi organization | 144 | 3 | 0.02 | 1.00E-06 |
| GO:0044829 | positive regulation by host of viral genome replication | 13 | 2 | 0 | 1.70E-06 |
| GO:0070309 | lens fiber cell morphogenesis | 13 | 2 | 0 | 1.70E-06 |
| GO:0002089 | lens morphogenesis in camera-type eye | 15 | 2 | 0 | 2.20E-06 |
| GO:0070307 | lens fiber cell development | 16 | 2 | 0 | 2.60E-06 |
| GO:0090110 | cargo loading into COPII-coated vesicle | 16 | 2 | 0 | 2.60E-06 |
| GO:0044794 | positive regulation by host of viral process | 18 | 2 | 0 | 3.30E-06 |
| GO:0044827 | modulation by host of viral genome replication | 21 | 2 | 0 | 4.50E-06 |
| GO:0070306 | lens fiber cell differentiation | 23 | 2 | 0 | 5.40E-06 |
| GO:0033363 | secretory granule organization | 24 | 2 | 0 | 5.90E-06 |
| GO:0035459 | cargo loading into vesicle | 24 | 2 | 0 | 5.90E-06 |
| GO:0044788 | modulation by host of viral process | 26 | 2 | 0 | 6.90E-06 |
| GO:0034389 | lipid particle organization | 27 | 2 | 0 | 7.50E-06 |
| GO:0060628 | regulation of ER to Golgi vesicle-mediated transport | 27 | 2 | 0 | 7.50E-06 |
| GO:0045070 | positive regulation of viral genome replication | 44 | 2 | 0.01 | 2.00E-05 |
| GO:0002088 | lens development in camera-type eye | 45 | 2 | 0.01 | 2.10E-05 |
| **Cellular Components** | | | | | |
| GO:0033116 | endoplasmic reticulum-Golgi intermediate compartment membrane | 42 | 2 | 0.01 | 1.90E-05 |
| GO:0030173 | integral component of Golgi membrane | 60 | 2 | 0.01 | 3.90E-05 |
| GO:0031228 | intrinsic component of Golgi membrane | 61 | 2 | 0.01 | 4.00E-05 |
| GO:0005793 | endoplasmic reticulum-Golgi intermediate compartment | 105 | 2 | 0.02 | 0.00012 |
| GO:0098791 | Golgi subcompartment | 849 | 3 | 0.14 | 0.00021 |
| GO:0044431 | Golgi apparatus part | 898 | 3 | 0.15 | 0.00025 |
| GO:0031301 | integral component of organelle membrane | 220 | 2 | 0.04 | 0.00052 |
| GO:0031300 | intrinsic component of organelle membrane | 248 | 2 | 0.04 | 0.00066 |
| GO:0031965 | nuclear membrane | 280 | 2 | 0.05 | 0.00084 |
| GO:1990730 | VCP-NSFL1C complex | 6 | 1 | 0 | 0.00099 |
| GO:0005794 | Golgi apparatus | 1650 | 3 | 0.27 | 0.00149 |
| GO:0005635 | nuclear envelope | 476 | 2 | 0.08 | 0.0024 |
| GO:0031616 | spindle pole centrosome | 18 | 1 | 0 | 0.00297 |
| GO:0031984 | organelle subcompartment | 2204 | 3 | 0.36 | 0.00346 |
| GO:0000139 | Golgi membrane | 678 | 2 | 0.11 | 0.0048 |
| GO:0045111 | intermediate filament cytoskeleton | 64 | 1 | 0.01 | 0.01054 |
| GO:0005789 | endoplasmic reticulum membrane | 1237 | 2 | 0.2 | 0.0154 |
| GO:0098827 | endoplasmic reticulum subcompartment | 1243 | 2 | 0.21 | 0.01554 |
| GO:0042175 | nuclear outer membrane-endoplasmic reticulum membrane network | 1256 | 2 | 0.21 | 0.01585 |
| GO:0044432 | endoplasmic reticulum part | 1480 | 2 | 0.24 | 0.02168 |
| **Molecular Functions** | | | | | |
| GO:0017137 | Rab GTPase binding | 150 | 2 | 0.03 | 0.00028 |
| GO:0019899 | enzyme binding | 2585 | 4 | 0.45 | 0.00042 |
| GO:0005096 | GTPase activator activity | 232 | 2 | 0.04 | 0.00066 |
| GO:0030695 | GTPase regulator activity | 247 | 2 | 0.04 | 0.00075 |
| GO:0060589 | nucleoside-triphosphatase regulator activity | 302 | 2 | 0.05 | 0.00112 |
| GO:0017016 | Ras GTPase binding | 395 | 2 | 0.07 | 0.0019 |
| GO:0031267 | small GTPase binding | 406 | 2 | 0.07 | 0.002 |
| GO:0008047 | enzyme activator activity | 488 | 2 | 0.08 | 0.00288 |
| GO:0051020 | GTPase binding | 561 | 2 | 0.1 | 0.00378 |
| GO:0009931 | calcium-dependent protein serine/threonine kinase activity | 24 | 1 | 0 | 0.00416 |
| GO:0010857 | calcium-dependent protein kinase activity | 24 | 1 | 0 | 0.00416 |
| GO:0051019 | mitogen-activated protein kinase binding | 27 | 1 | 0 | 0.00468 |
| GO:0004300 | enoyl-CoA hydratase activity | 41 | 1 | 0.01 | 0.0071 |
| GO:0003860 | 3-hydroxyisobutyryl-CoA hydrolase activity | 44 | 1 | 0.01 | 0.00762 |
| GO:0051117 | ATPase binding | 61 | 1 | 0.01 | 0.01055 |
| GO:0030234 | enzyme regulator activity | 1000 | 2 | 0.17 | 0.01163 |
| GO:0016289 | CoA hydrolase activity | 74 | 1 | 0.01 | 0.01279 |
| GO:0016790 | thiolester hydrolase activity | 104 | 1 | 0.02 | 0.01793 |
| GO:0098772 | molecular function regulator | 1347 | 2 | 0.23 | 0.02054 |
| GO:0043130 | ubiquitin binding | 124 | 1 | 0.02 | 0.02135 |

**Table 2:** Sponges without competitors vs sponges with soft coral *Zoanthus sansibaricus* as the competitor – down-regulated genes

| GO.ID | Term | Annotated | Significant | Expected | classicFisher |
|---|---|---|---|---|---|
| | **Biological Processes** | | | | |
| GO:0006119 | oxidative phosphorylation | 915 | 10 | 0.66 | 2.60E-10 |
| GO:0009126 | purine nucleoside monophosphate metabolic process | 1725 | 12 | 1.24 | 3.80E-10 |
| GO:0009167 | purine ribonucleoside monophosphate metabolic process | 1725 | 12 | 1.24 | 3.80E-10 |
| GO:0009161 | ribonucleoside monophosphate metabolic process | 1781 | 12 | 1.28 | 5.50E-10 |
| GO:0009123 | nucleoside monophosphate metabolic process | 1812 | 12 | 1.3 | 6.70E-10 |
| GO:0009150 | purine ribonucleotide metabolic process | 1924 | 12 | 1.38 | 1.30E-09 |
| GO:0006163 | purine nucleotide metabolic process | 1965 | 12 | 1.41 | 1.70E-09 |
| GO:0009259 | ribonucleotide metabolic process | 1989 | 12 | 1.43 | 1.90E-09 |
| GO:0019693 | ribose phosphate metabolic process | 2056 | 12 | 1.48 | 2.80E-09 |
| GO:0072521 | purine-containing compound metabolic process | 2069 | 12 | 1.49 | 3.10E-09 |
| GO:0046034 | ATP metabolic process | 1621 | 11 | 1.17 | 3.70E-09 |
| GO:0009205 | purine ribonucleoside triphosphate metabolic process | 1705 | 11 | 1.23 | 6.30E-09 |
| GO:0009199 | ribonucleoside triphosphate metabolic processs | 1730 | 11 | 1.24 | 7.30E-09 |
| GO:0009144 | purine nucleoside triphosphate metabolic process | 1732 | 11 | 1.25 | 7.40E-09 |
| GO:0045333 | cellular respiration | 1297 | 10 | 0.93 | 7.40E-09 |
| GO:0009141 | nucleoside triphosphate metabolic process | 1789 | 11 | 1.29 | 1.00E-08 |
| GO:0015980 | energy derivation by oxidation of organic compounds | 1383 | 10 | 1 | 1.40E-08 |
| GO:0017144 | drug metabolic process | 2962 | 13 | 2.13 | 1.40E-08 |
| GO:0009117 | nucleotide metabolic process | 2383 | 12 | 1.71 | 1.50E-08 |
| GO:0006753 | nucleoside phosphate metabolic process | 2390 | 12 | 1.72 | 1.60E-08 |
| | **Cellular Components** | | | | |
| GO:0005743 | mitochondrial inner membrane | 1803 | 14 | 1.49 | 1.60E-11 |
| GO:0019866 | organelle inner membrane | 1846 | 14 | 1.53 | 2.20E-11 |
| GO:0070469 | respiratory chain | 1132 | 12 | 0.94 | 2.40E-11 |
| GO:0031966 | mitochondrial membrane | 2144 | 14 | 1.77 | 1.60E-10 |
| GO:0044429 | mitochondrial part | 2666 | 15 | 2.2 | 2.00E-10 |
| GO:0005740 | mitochondrial envelope | 2227 | 14 | 1.84 | 2.70E-10 |
| GO:0005739 | mitochondrion | 3891 | 16 | 3.22 | 3.50E-09 |
| GO:0031967 | organelle envelope | 2832 | 14 | 2.34 | 6.40E-09 |
| GO:0070069 | cytochrome complex | 536 | 8 | 0.44 | 7.70E-09 |
| GO:0031975 | envelope | 2927 | 14 | 2.42 | 9.80E-09 |
| GO:0098803 | respiratory chain complex | 689 | 8 | 0.57 | 5.40E-08 |
| GO:0031090 | organelle membrane | 4640 | 14 | 3.84 | 3.30E-06 |
| GO:0045277 | respiratory chain complex IV | 360 | 5 | 0.3 | 1.00E-05 |
| GO:0016021 | integral component of membrane | 6836 | 15 | 5.65 | 6.30E-05 |
| GO:0098796 | membrane protein complex | 2377 | 9 | 1.97 | 7.20E-05 |
| GO:0031224 | intrinsic component of membrane | 6941 | 15 | 5.74 | 7.60E-05 |
| GO:0045275 | respiratory chain complex III | 178 | 3 | 0.15 | 0.00042 |
| GO:0043231 | intracellular membrane-bounded organelle | 15733 | 21 | 13.01 | 0.0009 |
| GO:0044425 | membrane part | 8863 | 15 | 7.33 | 0.00136 |
| GO:0043227 | membrane-bounded organelle | 16798 | 21 | 13.89 | 0.00272 |
| | **Molecular Function** | | | | |
| GO:0009055 | electron transfer activity | 1092 | 11 | 0.82 | 1.50E-10 |
| GO:0004129 | cytochrome-c oxidase activity | 548 | 8 | 0.41 | 4.60E-09 |
| GO:0015002 | heme-copper terminal oxidase activity | 548 | 8 | 0.41 | 4.60E-09 |
| GO:0016676 | oxidoreductase activity, acting on a heme group donors, oxygen as acceptor | 548 | 8 | 0.41 | 4.60E-09 |
| GO:0016675 | oxidoreductase activity, acting on a heme group donors | 552 | 8 | 0.42 | 4.90E-09 |
| GO:0015078 | proton transmembrane transporter activity | 977 | 9 | 0.74 | 2.30E-08 |
| GO:0015077 | monovalent inorganic cation transmembrane transporter activity | 1092 | 9 | 0.82 | 5.90E-08 |
| GO:0022890 | inorganic cation transmembrane transporter activity | 1285 | 9 | 0.97 | 2.30E-07 |
| GO:0015075 | ion transmembrane transporter activity | 1728 | 10 | 1.3 | 2.40E-07 |
| GO:0008324 | cation transmembrane transporter activity | 1387 | 9 | 1.04 | 4.50E-07 |
| GO:0015318 | inorganic molecular entity transmembrane transporter activity | 1576 | 9 | 1.19 | 1.30E-06 |
| GO:0016491 | oxidoreductase activity | 4106 | 13 | 3.09 | 2.10E-06 |
| GO:0022857 | transmembrane transporter activity | 2351 | 10 | 1.77 | 4.00E-06 |
| GO:0005215 | transporter activity | 2469 | 10 | 1.86 | 6.30E-06 |
| GO:0020037 | heme binding | 637 | 5 | 0.48 | 0.0001 |
| GO:0008121 | ubiquinol-cytochrome-c reductase activity | 157 | 3 | 0.12 | 0.00022 |
| GO:0016681 | oxidoreductase activity, acting on diphenols and related subbstances as donors | 157 | 3 | 0.12 | 0.00022 |
| GO:0016679 | oxidoreductase activity, acting on diphenols and related subbstances as donors | 177 | 3 | 0.13 | 0.00032 |
| GO:0046906 | tetrapyrrole binding | 888 | 5 | 0.67 | 0.00047 |
| GO:0055056 | D-glucose transmembrane transporter activity | 1 | 1 | 0 | 0.00075 |

**Table 3:** Sponges without competitors vs sponges with macroalgae *Dictyota ciliolata* as the competitor – up-regulated genes

| GO.ID | Term | Annotated | Significant | Expected | classicFisher |
|---|---|---|---|---|---|
| | **Biological Processes** | | | | |
| GO:0022900 | electron transport chain | 780 | 1 | 0.03 | 0.026 |
| GO:0006091 | generation of precursor metabolites and energy | 2125 | 1 | 0.07 | 0.07 |
| GO:0055114 | oxidation-reduction process | 3210 | 1 | 0.1 | 0.105 |
| GO:0044237 | cellular metabolic process | 20834 | 1 | 0.68 | 0.681 |
| GO:0008152 | metabolic process | 22651 | 1 | 0.74 | 0.741 |
| GO:0009987 | cellular process | 27066 | 1 | 0.89 | 0.885 |
| GO:0008150 | biological_process | 30575 | 1 | 1 | 1 |
| GO:0000002 | mitochondrial genome maintenance | 54 | 0 | 0 | 1 |
| GO:0000003 | reproduction | 1660 | 0 | 0.05 | 1 |
| GO:0000011 | vacuole inheritance | 6 | 0 | 0 | 1 |
| GO:0000012 | single strand break repair | 15 | 0 | 0 | 1 |
| GO:0000018 | regulation of DNA recombination | 141 | 0 | 0 | 1 |
| GO:0000019 | regulation of mitotic recombination | 12 | 0 | 0 | 1 |
| GO:0000022 | mitotic spindle elongation | 11 | 0 | 0 | 1 |
| GO:0000023 | maltose metabolic process | 1 | 0 | 0 | 1 |
| GO:0000027 | ribosomal large subunit assembly | 344 | 0 | 0.01 | 1 |
| GO:0000028 | ribosomal small subunit assembly | 281 | 0 | 0.01 | 1 |
| GO:0000032 | cell wall mannoprotein biosynthetic process | 1 | 0 | 0 | 1 |
| GO:0000038 | very long-chain fatty acid metabolic process | 63 | 0 | 0 | 1 |
| GO:0000041 | transition metal ion transport | 167 | 0 | 0.01 | 1 |
| | **Cellular Components** | | | | |
| GO:0042151 | nematocyst | 12 | 1 | 0 | 0.00079 |
| GO:0044448 | cell cortex part | 228 | 1 | 0.02 | 0.01502 |
| GO:0005938 | cell cortex | 499 | 1 | 0.03 | 0.03273 |
| GO:0099568 | cytoplasmic region | 804 | 1 | 0.05 | 0.05247 |
| GO:0070469 | respiratory chain | 1132 | 1 | 0.07 | 0.07347 |
| GO:0005743 | mitochondrial inner membrane | 1803 | 1 | 0.12 | 0.1157 |
| GO:0019866 | organelle inner membrane | 1846 | 1 | 0.12 | 0.11837 |
| GO:0031966 | mitochondrial membrane | 2144 | 1 | 0.14 | 0.13678 |
| GO:0005740 | mitochondrial envelope | 2227 | 1 | 0.15 | 0.14188 |
| GO:0044429 | mitochondrial part | 2666 | 1 | 0.18 | 0.16856 |
| GO:0031967 | organelle envelope | 2832 | 1 | 0.19 | 0.17855 |
| GO:0031975 | envelope | 2927 | 1 | 0.19 | 0.18423 |
| GO:0005576 | extracellular region | 3101 | 1 | 0.21 | 0.19459 |
| GO:0005739 | mitochondrion | 3891 | 1 | 0.26 | 0.2408 |
| GO:0031090 | organelle membrane | 4640 | 1 | 0.31 | 0.28336 |
| GO:0044444 | cytoplasmic part | 17683 | 2 | 1.17 | 0.34198 |
| GO:0071944 | cell periphery | 5712 | 1 | 0.38 | 0.34212 |
| GO:0016021 | integral component of membrane | 6836 | 1 | 0.45 | 0.40104 |
| GO:0031224 | intrinsic component of membrane | 6941 | 1 | 0.46 | 0.40641 |
| GO:0043229 | intracellular organelle | 21310 | 2 | 1.41 | 0.49665 |
| | **Molecular Functions** | | | | |
| GO:0005507 | copper ion binding | 272 | 1 | 0.01 | 0.0079 |
| GO:0004129 | cytochrome-c oxidase activity | 548 | 1 | 0.02 | 0.0159 |
| GO:0015002 | heme-copper terminal oxidase activity | 548 | 1 | 0.02 | 0.0159 |
| GO:0016676 | oxidoreductase activity, acting on a heme group donors, oxygen as receptor | 548 | 1 | 0.02 | 0.0159 |
| GO:0016675 | oxidoreductase activity, acting on a heme group donors | 552 | 1 | 0.02 | 0.016 |
| GO:0015078 | proton transmembrane transporter activity | 977 | 1 | 0.03 | 0.0283 |
| GO:0009055 | electron transfer activity | 1092 | 1 | 0.03 | 0.0316 |
| GO:0015077 | monovalent inorganic cation transmembrane transporter activity | 1092 | 1 | 0.03 | 0.0316 |
| GO:0022890 | inorganic cation transmembrane transporter activity | 1285 | 1 | 0.04 | 0.0372 |
| GO:0008324 | cation transmembrane transporter activity | 1387 | 1 | 0.04 | 0.0402 |
| GO:0015318 | inorganic molecular entity transmembrane transporter activity | 1576 | 1 | 0.05 | 0.0456 |
| GO:0015075 | ion transmembrane transporter activity | 1728 | 1 | 0.05 | 0.05 |
| GO:0022857 | transmembrane transporter activity | 2351 | 1 | 0.07 | 0.0681 |
| GO:0005215 | transporter activity | 2469 | 1 | 0.07 | 0.0715 |
| GO:0046914 | transition metal ion binding | 2511 | 1 | 0.07 | 0.0727 |
| GO:0016491 | oxidoreductase activity | 4106 | 1 | 0.12 | 0.1189 |
| GO:0046872 | metal ion binding | 8427 | 1 | 0.24 | 0.244 |
| GO:0043169 | cation binding | 8498 | 1 | 0.25 | 0.2461 |
| GO:0043167 | ion binding | 15588 | 1 | 0.45 | 0.4514 |
| GO:0003824 | catalytic activity | 18229 | 1 | 0.53 | 0.5278 |

**Table 4:** Sponges without competitors vs sponges with macroalgae *Dictyota ciliolata* as the competitor – down-regulated genes

| GO.ID | Term | Annotated | Significant | Expected | classicFisher |
|---|---|---|---|---|---|
| | **Biological Processes** | | | | |
| GO:0009060 | aerobic respiration | 744 | 5 | 0.27 | 3.40E-06 |
| GO:0006119 | oxidative phosphorylation | 915 | 5 | 0.33 | 9.40E-06 |
| GO:0045333 | cellular respiration | 1297 | 5 | 0.47 | 5.10E-05 |
| GO:0015980 | energy derivation by oxidation of organic compounds | 1383 | 5 | 0.5 | 6.90E-05 |
| GO:0046034 | ATP metabolic process | 1621 | 5 | 0.58 | 0.00015 |
| GO:0009205 | purine ribonucleoside triphosphate metabolic process | 1705 | 5 | 0.61 | 0.00019 |
| GO:0009126 | purine nucleoside monophosphate metabolic process | 1725 | 5 | 0.62 | 0.0002 |
| GO:0009167 | purine ribonucleoside monophosphate metabolic process | 1725 | 5 | 0.62 | 0.0002 |
| GO:0009199 | ribonucleoside triphosphate metabolic process | 1730 | 5 | 0.62 | 0.0002 |
| GO:0009144 | purine nucleoside triphosphate metabolic process | 1732 | 5 | 0.62 | 0.0002 |
| GO:0009161 | ribonucleoside monophosphate metabolic process | 1781 | 5 | 0.64 | 0.00023 |
| GO:0009141 | nucleoside triphosphate metabolic process | 1789 | 5 | 0.64 | 0.00023 |
| GO:0061014 | positive regulation of mRNA catabolic process | 65 | 2 | 0.02 | 0.00024 |
| GO:0017144 | drug metabolic process | 2962 | 6 | 1.07 | 0.00025 |
| GO:0009123 | nucleoside monophosphate metabolic process | 1812 | 5 | 0.65 | 0.00025 |
| GO:0009150 | purine ribonucleotide metabolic process | 1924 | 5 | 0.69 | 0.00033 |
| GO:0006163 | purine nucleotide metabolic process | 1965 | 5 | 0.71 | 0.00036 |
| GO:0009259 | ribonucleotide metabolic process | 1989 | 5 | 0.72 | 0.00038 |
| GO:0019693 | ribose phosphate metabolic process | 2056 | 5 | 0.74 | 0.00045 |
| GO:1903313 | positive regulation of mRNA metabolic pr... | 89 | 2 | 0.03 | 0.00045 |
| | **Cellular Components** | | | | |
| GO:0045277 | respiratory chain complex IV | 360 | 5 | 0.15 | 2.80E-07 |
| GO:0070069 | cytochrome complex | 536 | 5 | 0.23 | 2.00E-06 |
| GO:0070469 | respiratory chain | 1132 | 6 | 0.49 | 3.70E-06 |
| GO:0098803 | respiratory chain complex | 689 | 5 | 0.3 | 6.70E-06 |
| GO:0044429 | mitochondrial part | 2666 | 7 | 1.15 | 4.40E-05 |
| GO:0005743 | mitochondrial inner membrane | 1803 | 6 | 0.78 | 5.30E-05 |
| GO:0019866 | organelle inner membrane | 1846 | 6 | 0.79 | 6.10E-05 |
| GO:0031966 | mitochondrial membrane | 2144 | 6 | 0.92 | 0.00014 |
| GO:0005740 | mitochondrial envelope | 2227 | 6 | 0.96 | 0.00017 |
| GO:0098796 | membrane protein complex | 2377 | 6 | 1.02 | 0.00025 |
| GO:0005739 | mitochondrion | 3891 | 7 | 1.67 | 0.00049 |
| GO:0031967 | organelle envelope | 2832 | 6 | 1.22 | 0.00064 |
| GO:0031975 | envelope | 2927 | 6 | 1.26 | 0.00077 |
| GO:0031265 | CD95 death-inducing signaling complex | 3 | 1 | 0 | 0.00129 |
| GO:0030690 | Noc1p-Noc2p complex | 4 | 1 | 0 | 0.00172 |
| GO:0097342 | ripoptosome | 4 | 1 | 0 | 0.00172 |
| GO:0031264 | death-inducing signaling complex | 6 | 1 | 0 | 0.00258 |
| GO:0043231 | intracellular membrane-bounded organelle | 15733 | 12 | 6.76 | 0.00266 |
| GO:0030689 | Noc complex | 8 | 1 | 0 | 0.00343 |
| GO:0043227 | membrane-bounded organelle | 16798 | 12 | 7.22 | 0.00546 |
| | **Molecular Functions** | | | | |
| GO:0004129 | cytochrome-c oxidase activity | 548 | 6 | 0.21 | 2.40E-08 |
| GO:0015002 | heme-copper terminal oxidase activity | 548 | 6 | 0.21 | 2.40E-08 |
| GO:0016676 | oxidoreductase activity, acting on a heme group donors, oxygen as acceptor | 548 | 6 | 0.21 | 2.40E-08 |
| GO:0016675 | oxidoreductase activity, acting on a heme group donors | 552 | 6 | 0.21 | 2.50E-08 |
| GO:0009055 | electron transfer activity | 1092 | 7 | 0.41 | 4.50E-08 |
| GO:0015078 | proton transmembrane transporter activity | 977 | 6 | 0.37 | 7.30E-07 |
| GO:0015077 | monovalent inorganic cation transmembrane transporter activity | 1092 | 6 | 0.41 | 1.40E-06 |
| GO:0020037 | heme binding | 637 | 5 | 0.24 | 2.40E-06 |
| GO:0022890 | inorganic cation transmembrane transporter activity | 1285 | 6 | 0.48 | 3.60E-06 |
| GO:0008324 | cation transmembrane transporter activity | 1387 | 6 | 0.52 | 5.60E-06 |
| GO:0015318 | inorganic molecular entity transmembrane transporter activity | 1576 | 6 | 0.59 | 1.20E-05 |
| GO:0046906 | tetrapyrrole binding | 888 | 5 | 0.33 | 1.20E-05 |
| GO:0015075 | ion transmembrane transporter activity | 1728 | 6 | 0.65 | 2.00E-05 |
| GO:0022857 | transmembrane transporter activity | 2351 | 6 | 0.88 | 0.00011 |
| GO:0005215 | transporter activity | 2469 | 6 | 0.93 | 0.00015 |
| GO:0016491 | oxidoreductase activity | 4106 | 7 | 1.55 | 0.0003 |
| GO:0048037 | cofactor binding | 2878 | 6 | 1.08 | 0.00034 |
| GO:0035877 | death effector domain binding | 3 | 1 | 0 | 0.00113 |
| GO:0097199 | cysteine-type endopeptidase activity inv... | 8 | 1 | 0 | 0.00301 |
| GO:0005123 | death receptor binding | 9 | 1 | 0 | 0.00338 |

**Table 5:** Sponges with macroalgae *Dictyota ciliolata* as the competitor vs sponges with soft coral *Zoanthus sansibaricus* as the competitor – up-regulated genes

| GO.ID | Term | Annotated | Significant | Expected | classicFisher |
|---|---|---|---|---|---|
| **Biological Processes** | | | | | |
| GO:0009060 | aerobic respiration | 744 | 1 | 0.05 | 0.048 |
| GO:0006119 | oxidative phosphorylation | 915 | 1 | 0.06 | 0.059 |
| GO:0045333 | cellular respiration | 1297 | 1 | 0.08 | 0.083 |
| GO:0015980 | energy derivation by oxidation of organic compounds | 1383 | 1 | 0.09 | 0.088 |
| GO:0046034 | ATP metabolic process | 1621 | 1 | 0.11 | 0.103 |
| GO:0009205 | purine ribonucleoside triphosphate metabolic process | 1705 | 1 | 0.11 | 0.108 |
| GO:0009126 | purine nucleoside monophosphate metabolic process | 1725 | 1 | 0.11 | 0.11 |
| GO:0009167 | purine ribonucleoside monophosphate metabolic process | 1725 | 1 | 0.11 | 0.11 |
| GO:0009199 | ribonucleoside triphosphate metabolic process | 1730 | 1 | 0.11 | 0.11 |
| GO:0009144 | purine nucleoside triphosphate metabolic process | 1732 | 1 | 0.11 | 0.11 |
| GO:0009161 | ribonucleoside monophosphate metabolic process | 1781 | 1 | 0.12 | 0.113 |
| GO:0009141 | nucleoside triphosphate metabolic process | 1789 | 1 | 0.12 | 0.114 |
| GO:0009123 | nucleoside monophosphate metabolic process | 1812 | 1 | 0.12 | 0.115 |
| GO:0009150 | purine ribonucleotide metabolic process | 1924 | 1 | 0.13 | 0.122 |
| GO:0006163 | purine nucleotide metabolic process | 1965 | 1 | 0.13 | 0.124 |
| GO:0009259 | ribonucleotide metabolic process | 1989 | 1 | 0.13 | 0.126 |
| GO:0019693 | ribose phosphate metabolic process | 2056 | 1 | 0.13 | 0.13 |
| GO:0072521 | purine-containing compound metabolic process | 2069 | 1 | 0.14 | 0.131 |
| GO:0006091 | generation of precursor metabolites and energy | 2125 | 1 | 0.14 | 0.134 |
| GO:0009117 | nucleotide metabolic process | 2383 | 1 | 0.16 | 0.15 |
| **Cellular Components** | | | | | |
| GO:0045277 | respiratory chain complex IV | 360 | 1 | 0.02 | 0.024 |
| GO:0070069 | cytochrome complex | 536 | 1 | 0.04 | 0.035 |
| GO:0098803 | respiratory chain complex | 689 | 1 | 0.05 | 0.045 |
| GO:0070469 | respiratory chain | 1132 | 1 | 0.07 | 0.073 |
| GO:0015935 | small ribosomal subunit | 1229 | 1 | 0.08 | 0.08 |
| GO:0005743 | mitochondrial inner membrane | 1803 | 1 | 0.12 | 0.116 |
| GO:0019866 | organelle inner membrane | 1846 | 1 | 0.12 | 0.118 |
| GO:0031966 | mitochondrial membrane | 2144 | 1 | 0.14 | 0.137 |
| GO:0005740 | mitochondrial envelope | 2227 | 1 | 0.15 | 0.142 |
| GO:0098796 | membrane protein complex | 2377 | 1 | 0.16 | 0.151 |
| GO:0032991 | protein-containing complex | 11984 | 2 | 0.79 | 0.157 |
| GO:0044391 | ribosomal subunit | 2665 | 1 | 0.18 | 0.169 |
| GO:0044429 | mitochondrial part | 2666 | 1 | 0.18 | 0.169 |
| GO:0031967 | organelle envelope | 2832 | 1 | 0.19 | 0.179 |
| GO:0031975 | envelope | 2927 | 1 | 0.19 | 0.184 |
| GO:0044446 | intracellular organelle part | 14596 | 2 | 0.97 | 0.233 |
| GO:0044422 | organelle part | 14816 | 2 | 0.98 | 0.24 |
| GO:0005739 | mitochondrion | 3891 | 1 | 0.26 | 0.241 |
| GO:0005840 | ribosome | 3921 | 1 | 0.26 | 0.243 |
| GO:0031090 | organelle membrane | 4640 | 1 | 0.31 | 0.283 |
| **Molecular Functions** | | | | | |
| GO:0004129 | cytochrome-c oxidase activity | 548 | 1 | 0.05 | 0.047 |
| GO:0015002 | heme-copper terminal oxidase activity | 548 | 1 | 0.05 | 0.047 |
| GO:0016676 | oxidoreductase activity, acting on a heme group of donors, oxygen as acceptor | 548 | 1 | 0.05 | 0.047 |
| GO:0016675 | oxidoreductase activity, acting on a heme group of donors | 552 | 1 | 0.05 | 0.047 |
| GO:0020037 | heme binding | 637 | 1 | 0.06 | 0.054 |
| GO:0046906 | tetrapyrrole binding | 888 | 1 | 0.08 | 0.075 |
| GO:0015078 | proton transmembrane transporter activity | 977 | 1 | 0.08 | 0.082 |
| GO:1901363 | heterocyclic compound binding | 15406 | 3 | 1.34 | 0.089 |
| GO:0097159 | organic cyclic compound binding | 15471 | 3 | 1.34 | 0.09 |
| GO:0009055 | electron transfer activity | 1092 | 1 | 0.09 | 0.092 |
| GO:0015077 | monovalent inorganic cation transmembrane transport activity | 1092 | 1 | 0.09 | 0.092 |
| GO:0003676 | nucleic acid binding | 6936 | 2 | 0.6 | 0.105 |
| GO:0022890 | inorganic cation transmembrane transport activity | 1285 | 1 | 0.11 | 0.108 |
| GO:0008324 | cation transmembrane transporter activity | 1387 | 1 | 0.12 | 0.116 |
| GO:0015318 | inorganic molecular entity transmembrane tansport activity | 1576 | 1 | 0.14 | 0.131 |
| GO:0015075 | ion transmembrane transporter activity | 1728 | 1 | 0.15 | 0.143 |
| GO:0022857 | transmembrane transporter activity | 2351 | 1 | 0.2 | 0.191 |
| GO:0005215 | transporter activity | 2469 | 1 | 0.21 | 0.2 |
| GO:0048037 | cofactor binding | 2878 | 1 | 0.25 | 0.23 |
| GO:0003735 | structural constituent of ribosome | 3667 | 1 | 0.32 | 0.286 |

**Table 6:** Sponges with macroalgae *Dictyota ciliolata* as the competitor vs sponges with soft coral *Zoanthus sansibaricus* as the competitor – down-regulated genes

| GO.ID | Term | Annotated | Significant | Expected | classicFisher |
|---|---|---|---|---|---|
| | **Biological Processes** | | | | |
| GO:0006412 | translation | 4738 | 61 | 12.55 | <1e-30 |
| GO:0043043 | peptide biosynthetic process | 4811 | 61 | 12.75 | <1e-30 |
| GO:0043604 | amide biosynthetic process | 4955 | 61 | 13.13 | <1e-30 |
| GO:0006518 | peptide metabolic process | 5038 | 61 | 13.35 | <1e-30 |
| GO:0043603 | cellular amide metabolic process | 5331 | 61 | 14.12 | 4.60E-30 |
| GO:0034641 | cellular nitrogen compound metabolic process | 13022 | 77 | 34.5 | 4.80E-24 |
| GO:1901566 | organonitrogen compound biosynthetic process | 7625 | 63 | 20.2 | 2.50E-23 |
| GO:0010467 | gene expression | 8833 | 64 | 23.4 | 1.20E-20 |
| GO:0034645 | cellular macromolecule biosynthetic process | 8207 | 61 | 21.74 | 1.30E-19 |
| GO:0009059 | macromolecule biosynthetic process | 8285 | 61 | 21.95 | 2.20E-19 |
| GO:0044271 | cellular nitrogen compound biosynthetic process | 9028 | 63 | 23.92 | 3.60E-19 |
| GO:1901564 | organonitrogen compound metabolic process | 15532 | 74 | 41.15 | 4.30E-15 |
| GO:0044267 | cellular protein metabolic process | 10588 | 61 | 28.05 | 8.60E-14 |
| GO:0044249 | cellular biosynthetic process | 11341 | 63 | 30.04 | 8.80E-14 |
| GO:1901576 | organic substance biosynthetic process | 11618 | 63 | 30.78 | 3.10E-13 |
| GO:0009058 | biosynthetic process | 11774 | 63 | 31.19 | 6.30E-13 |
| GO:0006807 | nitrogen compound metabolic process | 19066 | 77 | 50.51 | 5.60E-12 |
| GO:0019538 | protein metabolic process | 11594 | 61 | 30.72 | 8.10E-12 |
| GO:0044237 | cellular metabolic process | 20834 | 79 | 55.19 | 2.30E-11 |
| GO:0008152 | metabolic process | 22651 | 81 | 60.01 | 2.70E-11 |
| | **Cellular Components** | | | | |
| GO:0005840 | ribosome | 3921 | 64 | 10.89 | <1e-30 |
| GO:1990904 | ribonucleoprotein complex | 5114 | 65 | 14.21 | <1e-30 |
| GO:0044391 | ribosomal subunit | 2665 | 44 | 7.4 | 1.20E-24 |
| GO:0032991 | protein-containing complex | 11984 | 77 | 33.29 | 1.40E-23 |
| GO:0022626 | cytosolic ribosome | 2288 | 40 | 6.36 | 5.80E-23 |
| GO:0044445 | cytosolic part | 2566 | 41 | 7.13 | 3.80E-22 |
| GO:0044444 | cytoplasmic part | 17683 | 83 | 49.12 | 1.50E-18 |
| GO:0043232 | intracellular non-membrane-bounded organelle | 9841 | 64 | 27.34 | 2.70E-16 |
| GO:0043228 | non-membrane-bounded organelle | 9847 | 64 | 27.35 | 2.80E-16 |
| GO:0015935 | small ribosomal subunit | 1229 | 22 | 3.41 | 1.80E-12 |
| GO:0015934 | large ribosomal subunit | 1473 | 23 | 4.09 | 8.00E-12 |
| GO:0022627 | cytosolic small ribosomal subunit | 1057 | 20 | 2.94 | 8.00E-12 |
| GO:0022625 | cytosolic large ribosomal subunit | 1236 | 21 | 3.43 | 1.60E-11 |
| GO:0043229 | intracellular organelle | 21310 | 82 | 59.2 | 1.10E-10 |
| GO:0043226 | organelle | 21709 | 82 | 60.31 | 4.50E-10 |
| GO:0005737 | cytoplasm | 23134 | 83 | 64.27 | 4.40E-09 |
| GO:0005829 | cytosol | 7614 | 43 | 21.15 | 2.70E-07 |
| GO:0070469 | respiratory chain | 1132 | 15 | 3.14 | 4.80E-07 |
| GO:0044446 | intracellular organelle part | 14596 | 63 | 40.55 | 5.20E-07 |
| GO:0044424 | intracellular part | 25655 | 84 | 71.27 | 9.90E-07 |
| | **Molecular Functions** | | | | |
| GO:0003735 | structural constituent of ribosome | 3667 | 61 | 8.92 | <1e-30 |
| GO:0005198 | structural molecule activity | 4840 | 61 | 11.77 | <1e-30 |
| GO:0003723 | RNA binding | 4463 | 29 | 10.86 | 3.20E-07 |
| GO:0004129 | cytochrome-c oxidase activity | 548 | 9 | 1.33 | 7.70E-06 |
| GO:0015002 | heme-copper terminal oxidase activity | 548 | 9 | 1.33 | 7.70E-06 |
| GO:0016676 | oxidoreductase activity, acting on a heme group of donors, oxygen as acceptor | 548 | 9 | 1.33 | 7.70E-06 |
| GO:0016675 | oxidoreductase activity, acting on a heme group of donors | 552 | 9 | 1.34 | 8.10E-06 |
| GO:0009055 | electron transfer activity | 1092 | 12 | 2.66 | 1.30E-05 |
| GO:0019843 | rRNA binding | 964 | 11 | 2.34 | 2.20E-05 |
| GO:0015078 | proton transmembrane transporter activity | 977 | 11 | 2.38 | 2.50E-05 |
| GO:0015077 | monovalent inorganic cation transmembrane transporter activity | 1092 | 11 | 2.66 | 6.80E-05 |
| GO:0022890 | inorganic cation transmembrane transporter activity | 1285 | 11 | 3.13 | 0.00028 |
| GO:0008324 | cation transmembrane transporter activity | 1387 | 11 | 3.37 | 0.00053 |
| GO:0020037 | heme binding | 637 | 7 | 1.55 | 0.00093 |
| GO:0003676 | nucleic acid binding | 6936 | 29 | 16.87 | 0.0014 |
| GO:0015318 | inorganic molecular entity transmembrane transporter activity | 1576 | 11 | 3.83 | 0.0015 |
| GO:0015075 | ion transmembrane transporter activity | 1728 | 11 | 4.2 | 0.00308 |
| GO:0046906 | tetrapyrrole binding | 888 | 7 | 2.16 | 0.00593 |
| GO:0008121 | ubiquinol-cytochrome-c reductase activity | 157 | 3 | 0.38 | 0.00671 |
| GO:0016681 | oxidoreductase activity, acting on diphenols and related substances | 157 | 3 | 0.38 | 0.00671 |
